# Using insects to detect, monitor and predict the distribution of *Xylella fastidiosa:* a case study in Corsica

**DOI:** 10.1101/241513

**Authors:** Astrid Cruaud, Anne-Alicia Gonzalez, Martin Godefroid, Sabine Nidelet, Jean-Claude Streito, Jean-Marc Thuillier, Jean-Pierre Rossi, Sylvain Santoni, Jean-Yves Rasplus

**Affiliations:** INRA, UMR1062 CBGP, F-34988 Montpellier, France; INRA, UMR1334 AGAP, F-34398 Montpellier, France

**Keywords:** qPCR, nested PCR, DNA extraction, plant-disease, insect vector, *Philaenus spumarius*

## Abstract

We sampled *ca* 2500 specimens of *Philaenus spumarius* throughout Corsica without *a priori* on the presence of symptoms on plants. We screened 448 specimens for the presence of *Xylella fastidiosa (Xf)* using qPCR and a custom nested PCR. qPCR appeared versatile and under-estimated the prevalence of *Xf*. Nested PCR showed that *Xf* was present in all populations. Molecular results were validated by prediction on the distribution of *Xf* made from tests conducted on plants, which shows the pertinence of using vectors in risk assessment studies. *Xf* was detected in tenerals and adults. Thus, *P. spumarius* could acquire *Xf* from its host plant, mostly *Cistus monspeliensis* in Corsica, which may act as reservoir for the next season. This contrasts with other observations and suggests that management strategies may have to be adapted on a case-by-case basis. At least two genetic entities and several variants of *Xf* not yet identified on plants were present in the insects, which suggests ancient introductions of *Xf* and a probable underestimation of the current diversity of the strains present in Corsica. Interestingly 6% of the specimens carried two subspecies. Studies are wanted to better characterize the strains present in Corsica and know how the disease was introduced, spread and why no sign of a potential epidemic was detected earlier. This study shows that, when sensitive enough methods are implemented, insects can be used to predict and better assess the exact distribution of *Xf*. Insects are indeed easy to collect, *Xf* multiply only in their foregut and does not become circulative, which facilitates its detection.

**Key message:**

- Insect vectors can be used to detect, monitor and predict the distribution of *Xylella fastidiosa*
- The widely used qPCR approach is not sensitive enough to detect low bacterial load
- Different strains/subspecies of *Xf* are widely distributed in Corsica which suggests old introduction(s)
- Strategies to manage *Xf* may need to be set up on a case-by-case basis
- There is an urgent need to take stock of the situation in Europe to avoid unnecessary economic pressure on certain geographical areas and agricultural sectors.

## INTRODUCTION

*Xylella fastidiosa (Xf)* (Xanthomonadaceae, Gammaproteobacteria) is a xylem-limited Gramn-egative bacterium that causes disease in important crops and ornamental plants, such as Pierce’s disease of grapevine, *Citrus* variegated chlorosis disease, phony peach disease, plum leaf scald as well as leaf scorch on almond or elm and *Quercus* (Retchless et al., 2014; Almeida and Nunney, 2015). *Xf* infects a large number of plants (more than 300 species from more than 60 plant families) (EFSA, 2015b, a). However, the different genetic lineages exhibit narrower host-plant ranges (Nunney et al., 2013). The disease is endemic and widespread on the American continent and its biology, ecology and epidemiology have been extensively studied in the last forty years (reviewed in (Redak et al., 2004a; Chatterjee et al., 2008; Janse and Obradovic, 2010; Purcell, 2013; Retchless et al., 2014; Almeida and Nunney, 2015).

In the last few years several subspecies of *Xf* have been detected in Europe (https://gd.eppo.int/taxon/XYLEFA/distribution,https://ec.europa.eu/food/plant/plant_health_biosecurity/legislation/emergency_measures/xylella-fastidiosa/latest-developments_en). An outbreak of *Xf pauca* was first identified in Apulia (southeastern Italy) in olive groves (Saponari et al., 2013a; Saponari et al., 2013b). *Xf multiplex* was then detected on *Polygala myrtifolia* in Corsica (Chauvel et al., 2015) and subsequently in continental southern France. Large-scale studies conducted in 2015 further revealed that *Xf pauca* and *Xf fastidiosa-sandyi* (ST76) as well as possible recombinants were also present in France (Denancé et al., 2017). Recently, *Xf multiplex* was detected in western parts of the Iberian peninsula (region of Alicante) and *Xf multiplex, Xf pauca* and *Xf fastidiosa* were detected in the Balearic Islands (Olmo et al., 2017). *Xf fastidiosa* was also detected in Germany on a potted *Nerium* oleander kept in glasshouse in winter (http://pflanzengesundheit.jki.bund.de/dokumente/upload/3a817_xylella-fastidiosa_pest-report.pdf) and several interception of infected coffee plants have been reported in Europe ((Jacques et al., 2016; Loconsole et al., 2016), https://gd.eppo.int/taxon/XYLEFA/distribution).

The bacterium is transmitted to plants by xylem-sap feeding leafhoppers (Hemiptera, Cicadomorpha) and members of several families are known to transmit the disease from plants to plants (Redak et al., 2004a; Redak et al., 2004b). In the Americas, Cicadellidae (Yang, 1994; Dellapé et al., 2016), spittlebugs (Cercopidae, Clastopteridae, Aphrophoridae) (Severin, 1949; Severin, 1950; Almeida et al., 2005; Krell et al., 2007), and cicadas (Cicadidae) (Paião et al., 2002; Krell et al., 2007) have been shown to efficiently transmit *Xf*. In Europe, few is known about the vectors that efficiently transmit the bacterium. So far, on the 119 potential vectors that feed on xylem sap (Chauvel et al., 2015) only *Philaenus spumarius* (Linnaeus, 1758), the meadow spittlebug, has been identified as an effective vector of *Xf* in southern Italy (Saponari et al., 2014; Cornara et al., 2016). Other studies are thus clearly needed to clarify whether other insects may play an important role in the epidemiology of *Xf*.

Usually, epidemiological survey of *Xf* is conducted on symptomatic plant material. Most frequently, the presence of the bacterium is assessed using qPCR targeting a small part of the gene *rimM* as designed in Harper et al. (2010 erratum 2013) (see Baldi & La Porta (2017) for a review of the available methods and their advantages and drawbacks). Then, if wanted, fragments of seven housekeeping genes are sequenced as defined in Yuan et al. (2010) and sequences are compared to a reference database (http://pubmlst.org/xfastidiosa/) to assign the strain to subspecies or detect recombinants (e.g. (Nunney et al., 2012a; Nunney et al., 2012b; European Plant Protection Organization, 2016; Jacques et al., 2016; Denancé et al., 2017). Large scale and unbiased survey of the disease requires exhaustive sampling of plants (both symptomatic and asymptomatic) in multiple habitats, which is fastidious. Furthermore, the heterogeneous distribution of the bacterium in the plant (EFSA, 2015a) as well as PCR inhibitors (e.g. polyphenols (Schrader et al., 2012)) may induce false-negative results. To the contrary, most insect vectors can be easily sampled through sweeping (among vectors only cicadas are relatively difficult to sample and may require acoustic tools to locate them).

Insects are also known to contain PCR inhibitors (Boncristiani et al., 2011; Shamim et al., 2014; R. Krugner USDA USA *pers. comm.*) but colonization of insects by *Xf* occurs in a non-circulative manner with bacterial colonies located in the foregut (Purcell & Finlay, 1979; observations on *Graphocephala atropunctata* (Signoret, 1854)). Thus, it is either possible to dissect the foregut of the insects or extract DNA from an entire specimen to make sure having access to the bacterium. Therefore, as suggested by the spy insect approach set up in buffer zones and symptom-less areas closed to contaminated olive groves in Italy (Yaseen et al., 2015; D’onghia, 2017; Yaseen et al., 2017), implementing massive survey to test whether or not insect populations in different ecosystems do carry *Xf* would efficiently complement studies on plants and improve the early detection and monitoring of the disease.

Here we propose to go one step further on this idea and provide a first assessment of the use of insects to detect, monitor or predict the distribution of *Xf* in Europe, using Corsica as a case study. In a first step, we propose to test the feasibility of a large screening of insect populations for the presence of *Xf* but also for the possible characterisation of the carried strains via PCR amplification and sequencing of the loci included in the MLST of *Xf*. We sampled 62 populations of *Philaenus spumarius* throughout Corsica (Fig. 1, Table S1) from early June (when there was still a mix of larvae and adults, Fig. S1) to late October (before the adults are presumably killed by winter). We then tested for the presence of *Xf* in a subset of 11 populations (448 specimens, Fig. 1, Table S1) using a qPCR approach and a nested PCR protocol designed for the purpose of the study. Indeed, targeting *Xf* using qPCR appeared inconclusive and did not allow assessing the genetic identity of the strains. In a second step, we compared the results of our molecular tests to the potential range of *Xf* as estimated using species distribution modelling based on presence / absence tests conducted on plants. In a third step, we collected occurrence data of *P. spumarius* throughout Europe and estimated its geographical range using species distribution modelling to discuss the interest of applying this spy insect approach to Europe.

**Figure 1.**
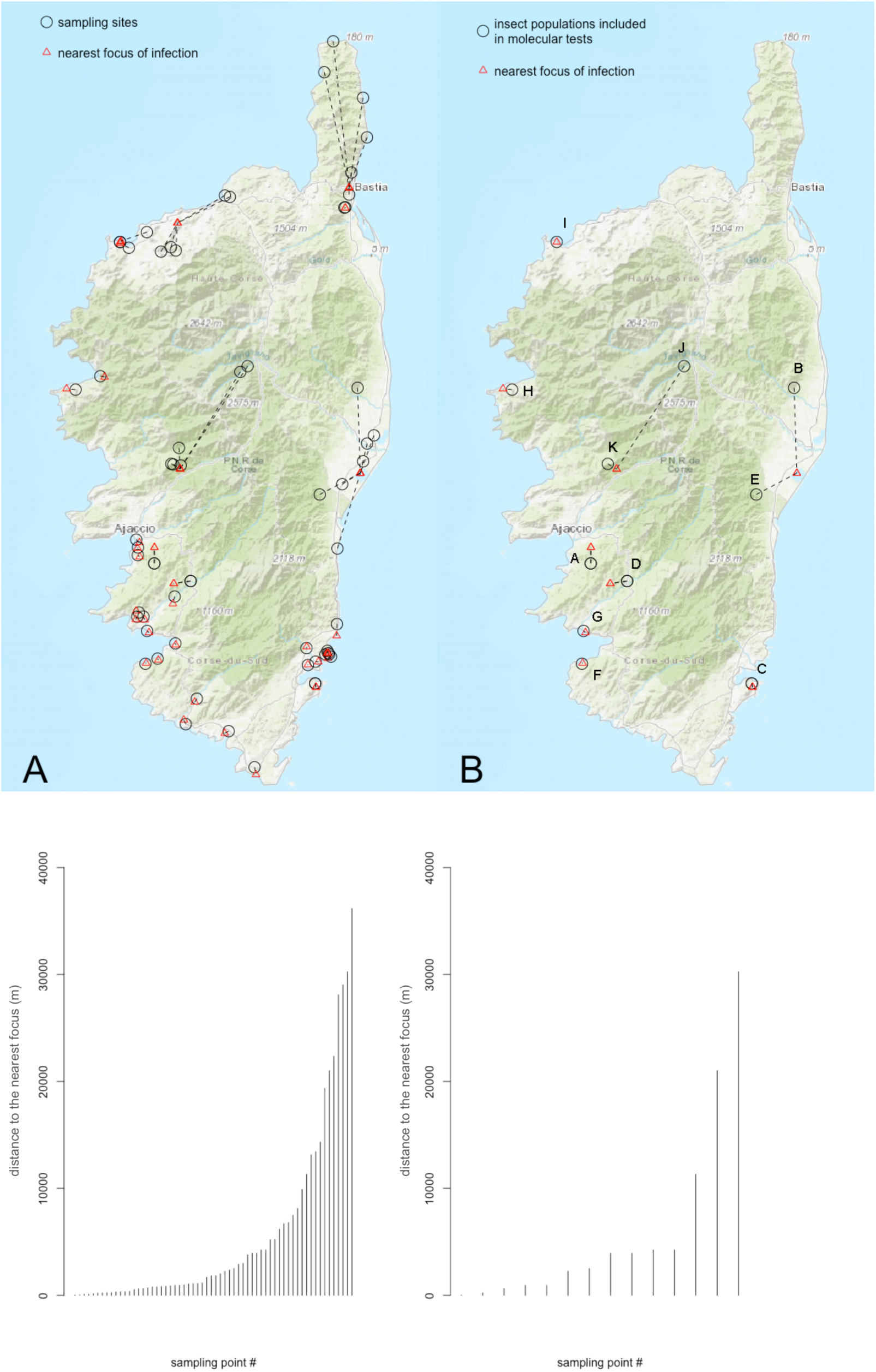
Map of the sampling sites and distance to the nearest focus of infection determined by molecular tests on plant material (national survey). A=all sampling sites, B=sampling sites where insect populations were tested for the presence of *Xf*. See Table S1 for more information on sampling sites.

## MATERIALS AND METHODS

### Sampling

We sampled adults of *Philaenus spumarius* in 62 localities in Corsica from early June to late October (Fig. 1, Table S1) by passing a sweep net through the vegetation using alternate backhand and forehand strokes. Specimens were collected in the net with an aspirator, killed on site with Ethyl Acetate and stored in 8mL vials containing 70% EtOH. Vials were stored in a freezer (-20°C) until DNA extraction. We mostly sampled in natural environments. The distance between sampling areas and the closest infested area (as identified by the national survey on plant material) ranged from ca. 20 m to 30000 m (Fig. 1). To optimize our sampling and test the feasibility of a large survey, we spent no more than 30 min sweeping in each locality. Specimens and plants on which they were successfully collected were identified to species. Molecular tests were conducted on a subset of 11 populations of *P. spumarius* (32 specimens per population, Fig. 1, Table S1). Three of these populations were sampled both in early June and late October to test for a possible seasonal variation of the prevalence of *Xf*. Other populations were sampled either in June (early / late) or October. A total of 448 specimens were screened for *Xf*.

### DNA extraction

It seems unfeasible to proceed to the dissection of the foregut of each specimen for a mass-survey. Moreover, dissection of insects can generate cross-contamination. Thus, we tried to improve each steps of the DNA extraction protocols classically used (Bextine et al., 2004; Brady et al., 2011) to reduce the impact of PCR inhibitors that may be contained in the insects (Boncristiani et al., 2011; Shamim et al., 2014) and increase yield in bacterial DNA. The complete protocol is available in Supplementary data (Appendix 1). Briefly, insects were placed in lysis buffer that contained PVP (Polyvinylpyrrolidone, to absorb polyphenols and polyamins thus preventing them from interacting with DNA which could inhibit PCR) and Sodium Bisulfite (to prevent oxydation of polyphenols, that, when oxidized covalently bind to DNA making it useless for further application). Insects were crushed using garnet crystals and ceramic beads coated with zirconium. Then, lysozyme was added to facilitate lysis of the bacteria. After 30min incubation, Proteinase K and extraction buffer that contained guanidium chlorure (to denature proteins and increase lysis of bacterial cells) and sodium bisulfite (antioxydant) were added to the mix. After one-hour incubation, deproteneisation using potassium acetate was performed. Finally, DNA extracts were purified using a KingFisher robot and Chemagic beads.

### Quantitative PCR (qPCR)

We used the method proposed by Harper et al. (2010 erratum 2013), which is listed as one of the official detection methods of *Xf* in plant material (European Plant Protection Organization, 2016) and recognized as the most sensitive method available to date for the detection of *Xf* in plants (Harper et al., 2010 erratum 2013; Baldi and La Porta, 2017). We followed recommendations by the ANSES (2015) and the EPPO (2016) but re-evaluated a cycle threshold using negative controls (ultrapure water and 2 *μ* g of phage lambda purified DNA) to better fit with our experimental conditions. To estimate the sensitivity of the qPCR approach, we used incremental dilution of an inactivated bacterial suspension of known concentration provided by B. Legendre (LSV, ANSES, Angers, France). Two replicates of qPCR were performed on each insect specimen.

### Nested PCR and Sequencing

Our attempt to amplify the seven loci included in the MLST of *Xf* using conventional PCR and primers / conditions described in the original protocol (Yuan et al., 2010, https://pubmlst.org/xfastidiosa/) were unsuccessful, probably because of the low amount of bacteria. We thus switched to a nested PCR approach. Sequences of the different alleles of each locus were downloaded from https://pubmlst.org/xfastidiosa/ (last access October 19th 2017) and aligned. Internal primers for each locus were designed from these alignments (Table S2) and primers were M13 tailed to simplify the sequencing reaction. We ensured that the nested PCR approach did not preclude discrimination among genetic entities by comparing maximum likelihood phylogenetic trees obtained from a concatenation of the 7 loci originally included in the MLST of *Xf* and their reduced sequences as included in the nested PCR scheme (Fig S2). Loci extracted from all genomes available on Genbank (last access October 19th 2017) were used as input. To test for the presence of *Xf* in the insects, we first targeted *holC*. When the amplification of *holC* was successful, a nested PCR to amplify the six other loci was attempted. *HolC* was first amplified using the primers listed in Yuan et al. (2010) and the mastermix and PCR conditions described in Tables S3 and S5. Five microliters of PCR product were then used to perform a nested PCR with the mastermix and PCR conditions described in Tables S3 and S5. For the six other loci, we first performed a triplex PCR (gltT/leuA/petC and cysG/malF/nuoL) using the primers listed in Yuan et al. (2010) and the mastermix and PCR conditions described in Tables S4 and S5. Five microliters of the PCR product were then used to perform a simplex nested PCR with the mastermix and PCR conditions described in Tables S4 and S5. The strict procedure implemented to avoid carry-over contamination is detailed in the Appendix 2 of the supplementary data file. To estimate the sensitivity of the nested PCR approach, we used incremental dilution of the same inactivated bacterial suspension as for qPCR. Sequencing of the PCR products was performed at AGAP on an Applied Biosystems 3500 Genetic Analyser. Allele assignation was performed using http://pubmlst.org/xfastidiosa/. Phylogenetic inferences were performed using raxmlHPC-PTHREADS-AVX (Stamatakis, 2014). Given that *α* and the proportion of invariable sites cannot be optimized independently from each other (Gu, 1995) and following Stamatakis’ personal recommendations (RAxML manual), a GTR + Γ model was applied to each gene region. We used a discrete gamma approximation (Yang, 1994) with four categories. GTRCAT approximation of models was used for ML boostrapping (Stamatakis, 2006) (1000 replicates). Resulting trees were visualised using Figtree (Rambaut, 2006). SplitsTree v.4.14.4 (Huson and Bryant, 2006) was used to build NeighborNet phylogenetic networks.

### Species distribution modelling of *Xf* at the Corsican scale

The potential distribution of *Xf* subsp. *multiplex* (ST6 & ST7) in Corsica was modelled using BIOCLIM (Busby, 1991) and DOMAIN (Carpenter et al., 1993). Methodology followed Godefroid et al. (submitted, see comments section below for preprint DOI). Data collected in France from 2015 to 2017 by the national survey on plant material completed by occurrences from the native area of the bacterium were used as input. Results were summarized as a suitability map averaging all model predictions.

### Species distribution modelling of *Phileanus spumarius* at the European scale

*Occurrence dataset:* A total of 1323 occurrences were used to model the distribution of *P. spumarius* in Europe (Fig. 6a). Off these, 471 originated from the GBIF database (GBIF.org (2017), *GBIF Home Page*. Available from: http://gbif.org [1rd November 2017]). The remaining 852 occurrences corresponded to our own observation records or were taken from the literature (List of references in Supplementary material, Appendix 3).

*Modelling framework:* We modelled the distribution of *P. spumarius* by means of Maxent, the most widely used species distribution model (Elith et al., 2006; Phillips et al., 2006). Maxent models the potential species distribution based on the principle of maximum entropy. It relates species occurrence records and background data with environmental descriptors to get insight into the environmental conditions that best reflect ecological requirements of the species. Species responses to environmental constraints are often complex, which implies using nonlinear functions (Elith et al., 2006). For that reason the parametrization of Maxent involves choosing among several transformations (referred to as feature classes or FCs) of original environmental descriptors (i.e. linear, quadratic, product, hinge and threshold: Phillips and Dudik, 2008). The parametrization of Maxent also involves a regularization multiplier (RM) introduced to reduce overfitting (Merow et al., 2013). Various authors have highlighted that the default settings of Maxent are not optimal in all situations and might lead to poorly performing models in certain cases (Merow et al., 2013; Shcheglovitova and Anderson, 2013; Radosavljevic and Anderson, 2014) and it is sensible to search for the best parameters given the dataset at hand (Radosavljevic and Anderson, 2014).

We built models with RM values ranging from 0.5 to 4 with increments of 0.5 and 6 FC combinations (L, LQ, H, LQH, LQHP, LQHPT with L=linear, Q=quadratic, H=hinge, P=product and T=threshold) using the R package ENMeval (Muscarella et al., 2014). This led to 48 different models. The models were fitted using all the available occurrence points and 10,000 randomly positioned background points. The optimal settings corresponded to the models giving the minimum AICc values (see Muscarella et al., 2014, for details). The resulting optimal FC and RM values were used to fit the final Maxent model based on a training dataset constituted by a random subset of 80% of the occurrences and 10,000 randomly positioned background points. The resulting Maxent model was then used to create a map of suitability scores (i.e. habitat suitability) across Western Europe (logistic output of Maxent: (Phillips and Dudik, 2008)). Suitability values were transformed into presence/absence by applying two thresholds a) the value at which the sum of the sensitivity (true positive rate) and specificity (true negative rate) is the highest and b) the highest value at which there is no omission (Liu et al., 2005b). These analyses were performed using the R package dismo (Hijmans et al., 2016).

We evaluated the performance of the model using the AUC metric (Fielding and Bell, 1997) and the Boyce index (Hirzel et al., 2006). The Boyce index is a reliable presence-only evaluation measure that varies between -1 and +1. Positive values indicate a model which predictions are consistent with the distribution of the presences in the evaluation dataset. The Boyce index was calculated using the R package ecospat (Broennimann et al., 2016).

*Environmental descriptors: bioclimatic variables:* Our modelling strategy relies on a set of bioclimatic descriptors hosted in the Worldclim database (Fick and Hijmans, 2017). Each variable is available in the form of a raster map and represent the average climate conditions for the period 1970–2000. We used raster layers of 2.5-minute spatial resolution, which corresponds to about 4.5 km at the equator. The choice of environmental descriptors to be involved is crucial and it is widely acknowledged that using reduced number of variables improves transferability and avoids model overparametrization (Peterson and Nakazawa, 2008; Jiménez-Valverde et al., 2011). We used the variables referred to as bio5, bio7 and bio19 corresponding to the maximum temperature of the warmest month, the temperature annual range and the precipitation of the coldest quarter (see details in Hijmans et al., 2005). Both temperature and precipitation were considered as proxy for the main environmental features constraining the insect distribution.

## RESULTS

### Detection of Xf in insect vectors using qPCR

Based on the results obtained with the negative controls, we fixed the cycle threshold (Ct) to 32.5. Thus, results were considered positive when Ct < 32.5 and an exponential amplification curve was observed, results were considered negative when Ct > 36 and results were considered undetermined when 32.5 < Ct < 36. Sensitivity tests on the inactivated bacterial suspension indicated that the signal was lost when i) less than 100 bacteria were present in the reaction mix or ii) less than 250 bacteria mixed with 2 μg of insect DNA were present in the reaction mix. The results of the two qPCR replicates were different for 43.8% of the insects (Fig. 2). A single positive qPCR was obtained for 2.7% of the insects collected in June. Four of the seven populations (sites A, B, D, J, Fig. 1) contained positive insects (from one to two positive insects per population). A single positive qPCR was obtained for 12.5% of the insects collected in October. Six of the seven populations (sites A, D, C, F, G, I) contained positive insects (from 3 to 7 positive insects per population). The two qPCR replicates were positive for 1.3% of the insects all collected in October from two populations (sites I: 2 positive insects; and G: 8 positive insects).

**Figure 2.**
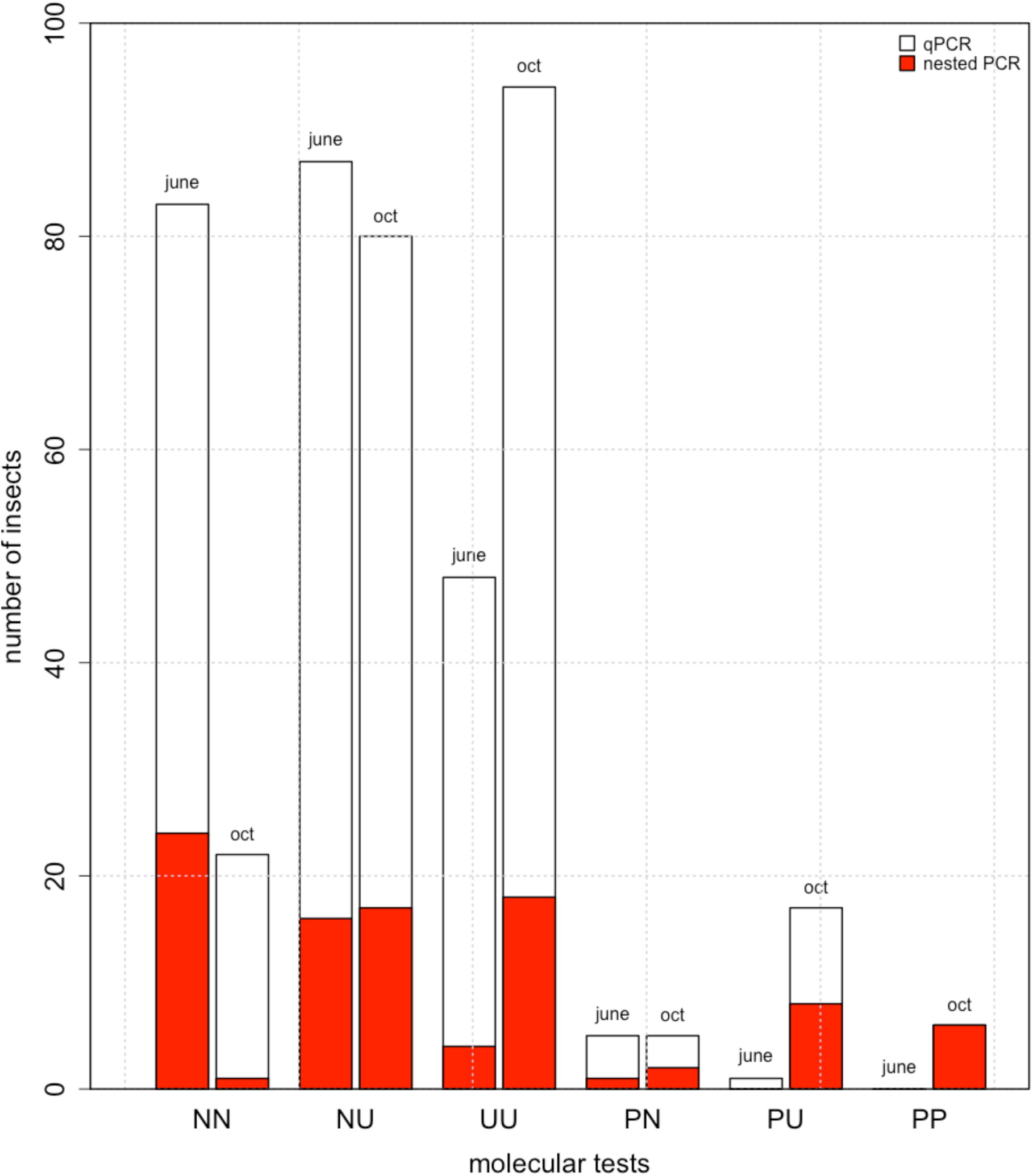
Results of the molecular tests performed on the insects (qPCR and nested PCR). The black histogram shows the distribution of the results of the qPCR replicates for the 448 insects. N = negative, P = positive, U = undetermined (see text for the Ct). Thus NN indicates that the two replicates were negative. The red histogram indicates that positive nested PCR on *holC* were obtained.

### Detection of Xf in insect vectors using nested PCR and sequencing

Sensitivity tests on the inactivated bacterial suspension indicated that the signal was lost when i) less than 5 bacteria were present in the reaction mix or ii) less than 50 bacteria mixed with 2 μg of insect DNA were present in the reaction mix. As compared with the qPCR approach, the nested PCR approach on *holC* revealed that *Xf* was present in all populations both in June and October with higher prevalence rates (Figs. 2 & 3). Positive nested PCR were always obtained when the results of the two qPCR replicates were positive, i.e. presumably from the insects with the highest bacterial load (collected in late October). However, positive nested PCR were also obtained when the results of the two qPCR replicates were negative (5.6% of the samples). The rate of false negative as compared to the nested PCR approach was 8.7% for the first replicate of qPCR and 10.5% for the second replicate. Notably, 11.2% (resp. 7.1%) of the qPCR that gave an undetermined result led to a positive nested PCR for the first replicate of qPCR (resp. the second replicate of qPCR). With the nested PCR approach, an average of 20.1% (23.2%) of the specimens were found positive to *Xf* in June (October). The prevalence of *Xf* in the different populations varied from 0.0% to 43.7 % in June and 12.5 - 34.4% in October. No significant seasonal variation of the presence of *Xf* was observed. Analysis of the sequences obtained for *holC* revealed that 56.7% of the insects tested positive for *Xf* carried allele *holC_3*, and 21.6% carried allele *holC_1* (Table 1, Fig. 4a). It is noteworthy that two yet undescribed variants of *holC_3* as well as two yet undescribed variants of *holC_1* were found in the screened populations. This result is interesting *per se* and indicates that the probability that our results are due to carry-over contamination is reduced. Interestingly for a few specimens (6% of the positive samples), double peaks were observed on the diagnostic sites for allele *holC_3* versus allele *holC_1*, which suggest that they may carry two subspecies of *Xf*.

**Table 1.** *HolC* alleles present in each population of *P. spumarius*.

|  | *holC 3* | *holC 1* |
| --- | --- | --- |
| Site A (June) | 1 | 1 |
| Site A (October) | 2 | 2 |
| Site C (June) | 5 + 1 variant | 0 |
| Site C (October) | 5 | 1 |
| Site D (June) | 0 | 0 |
| Site D (October) | 3 + 1 variant | 0 |
| Site B | 3 | 0 |
| Site E | 1 | 7 |
| Site J | 4 | 3 |
| Site K | 6 + 1 variant | 2 |
| Site F | 10 | 1 |
| Site G | 9 | 1 |
| Site H | 0 | 1 + 2 variants |
| Site I | 6 | 2 |

**Figure 3.**
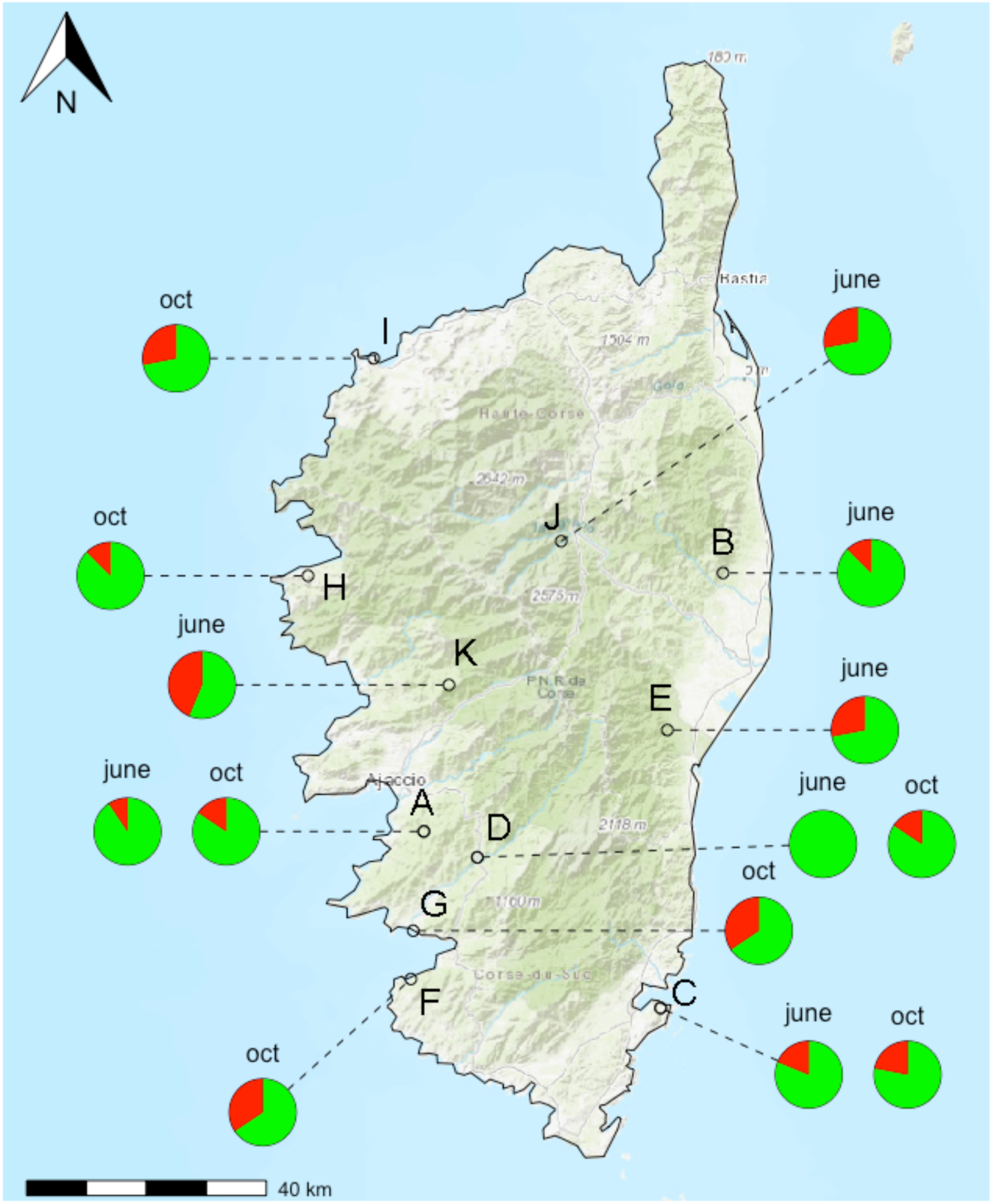
Prevalence of *Xf* in the different populations as revealed by the nested PCR approach targeting *holC*. Red (green) color indicates the percentage of insects tested positive (negative) for the presence of *Xf*. Tests were conducted on 32 specimens in each sample site.

**Figure 4.**
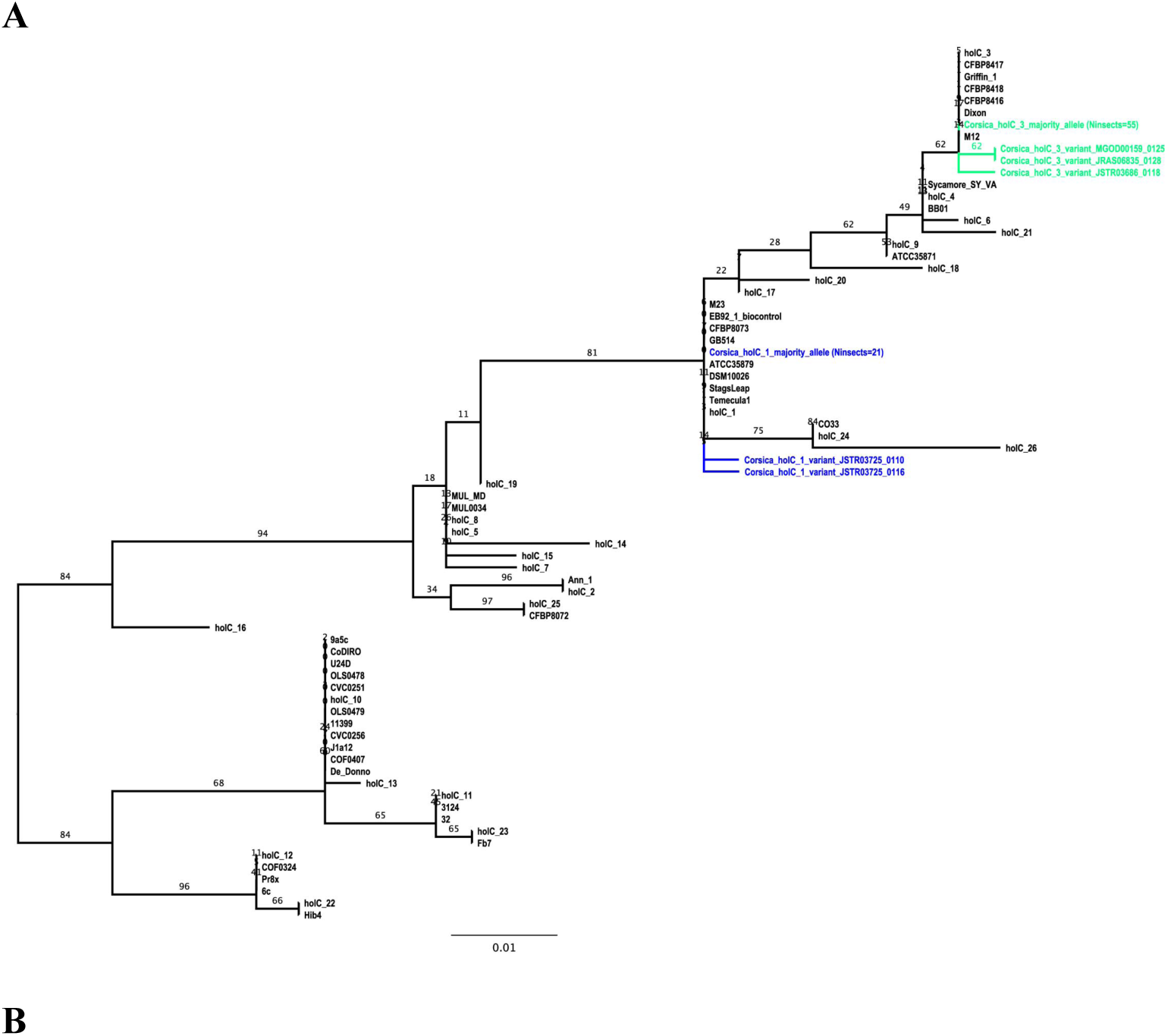
Position of the strains characterized in populations of insects in Corsica. A) RAxML tree inferred from the analysis of the reduced *holC* sequences targeted with the nested PCR approach (BP at nodes, 1000 replicates). Alleles present in https://pubmlst.org/xfastidiosa/ (last access October 19th 2017), genomes available on Genbank (last access October 19th 2017) and alleles obtained from insect samples are included in the analysis. B) NeighborNet phylogenetic network. Inferences are based on a concatenation of the reduced *leuA, petC, malF, cysG, holC, nuoL, gltT* targeted with the nested PCR approach. STs present in https://pubmlst.org/xfastidiosa/ as well as genomes available on Genbank are included in the analysis (last access October 19th 2017). Insect samples for which at least two loci could be sequenced (JSTR03697_0126 & JSTR03697_0129: 7 loci sequenced and MGOD00159_0113 & JRAS06849_0101: *holC* and *gltT* sequenced) are included in the network. Note: CO33 / ST72 (Giampetruzzi et al., 2015; Loconsole et al., 2016); ST76 (Loconsole et al., 2016); CFBP8073 / ST75 (Jacques et al., 2016) as well as ST79 (Denancé et al., 2017) are genetically related to isolates belonging to different subspecies of *Xf*.

Sequences for the seven loci of the MLST have been obtained for the specimens with the highest bacterial load (2 specimens collected in site G in October). The complete typing indicates that the carried strain was *Xf multiplex* ST7. Only partial typing (at most 2 loci) could be obtained for other specimens. Thus we could not conclude without doubt on the identity of the strains they carried. For two specimens that carried *holC_1*, one collected in late June in site K and one collected in October in site E, sequencing of *gltT* indicates that the carried allele was *gltT_1* and a variant of *gltT_1* respectively, which suggests that the subspecies *Xf fastidiosa* may be also present in Corsica, though this results needs to be confirmed by a complete typing. The NeighborNet network inferred from the concatenation of the reduced sequences of *leuA, petC, malF, cysG, holC, nuoL, gltT* targeted by the nested PCR approach and including reported STs, available genomes as well as Corsican strains characterized on more than one locus is presented in Figure 4b. All sequences have been deposited on Genbank (XXXX-XXX-upon acceptance-)

### Match between the molecular results and the predicted distribution of Xf multiplex ST6 & ST7

All sampling sites fall within the predicted distribution of the bacterium, which encompasses the entire island apart from mountainous regions in the centre (Fig. 5). Two sampling sites (stations J & K) fall near the edge of the predicted potential distribution area of *Xf*.

**Figure 5.**
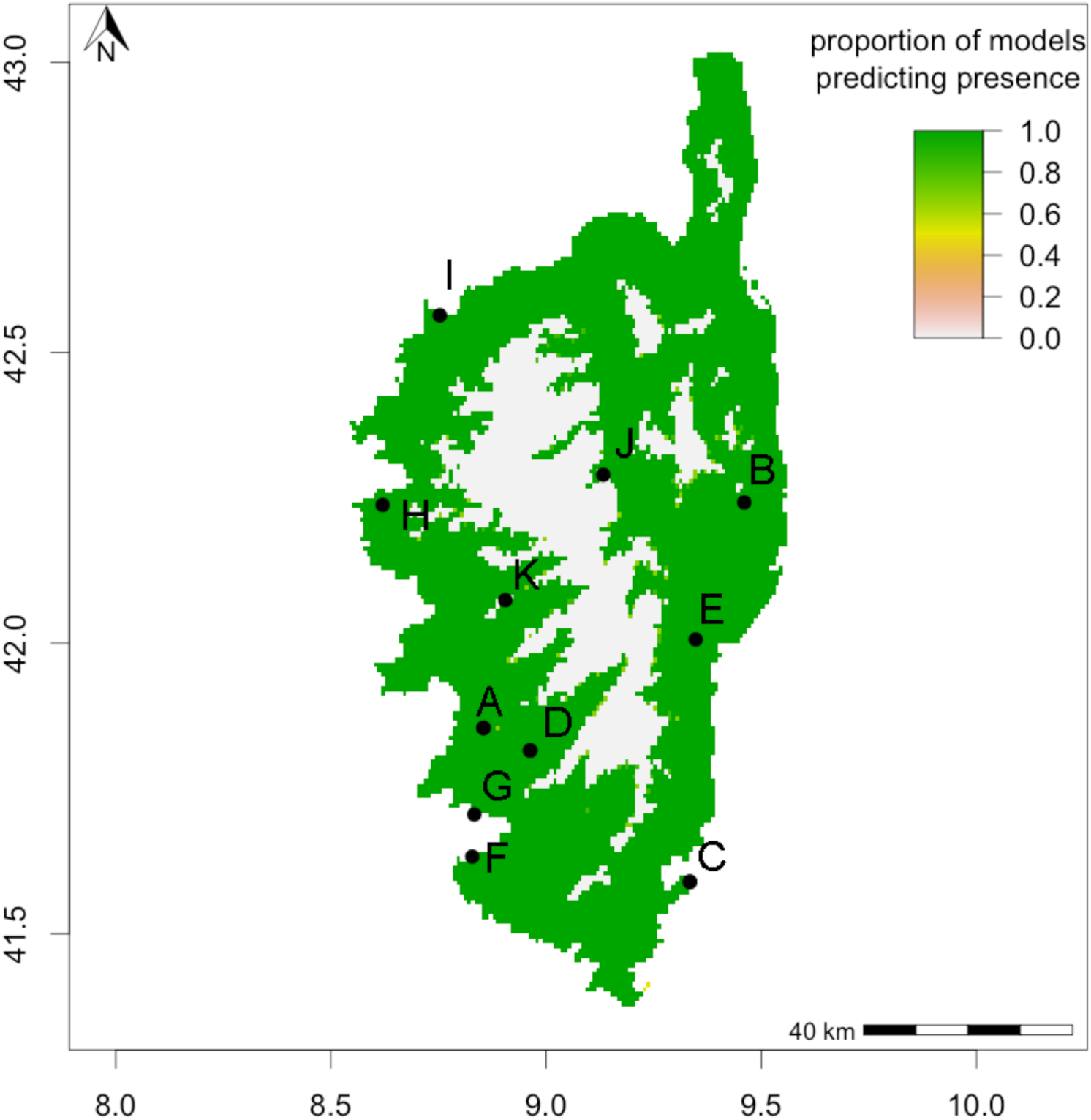
Potential geographical distribution of *Xf multiplex* (ST6/ST7) in Corsica. Suitability map averaging predictions by BIOCLIM and DOMAIN (from red/low to green/high suitability). Black dots indicate insect populations tested for the presence of *Xf*. See Godefroid et al. (submitted, see comments section below for preprint DOI) for details

### *Distribution of* Phileanus spumarius *at the European scale*

Occurrence data used in the study are presented in Figure 6a. The regularization multiplier of the Maxent model giving the minimum AICc values was 1.5 and it was associated to feature class combining linear, quadratic, hinge, product and threshold features (LQHPT). The value of the AUC of the Maxent model fitted using the latter optimal settings was 0.89 and the Boyce index was 0.986. Both metrics indicated that the Maxent model performed satisfactorily. Figure 6b displays the prediction of the Maxent model (logistic output) i.e. the habitat suitability for *P. spumarius* across the study area. Most of the Western Europe appeared to be associated to high habitat suitability values. The presence/absence maps derived from the conversion of habitat suitability using the threshold value maximizing the sensitivity and the specificity or the threshold for which there is no omission (Liu et al., 2005a) are given in Figure S3. In both cases nearly all the Western Europe appeared to host *P. spumarius*.

**Figure 6.**
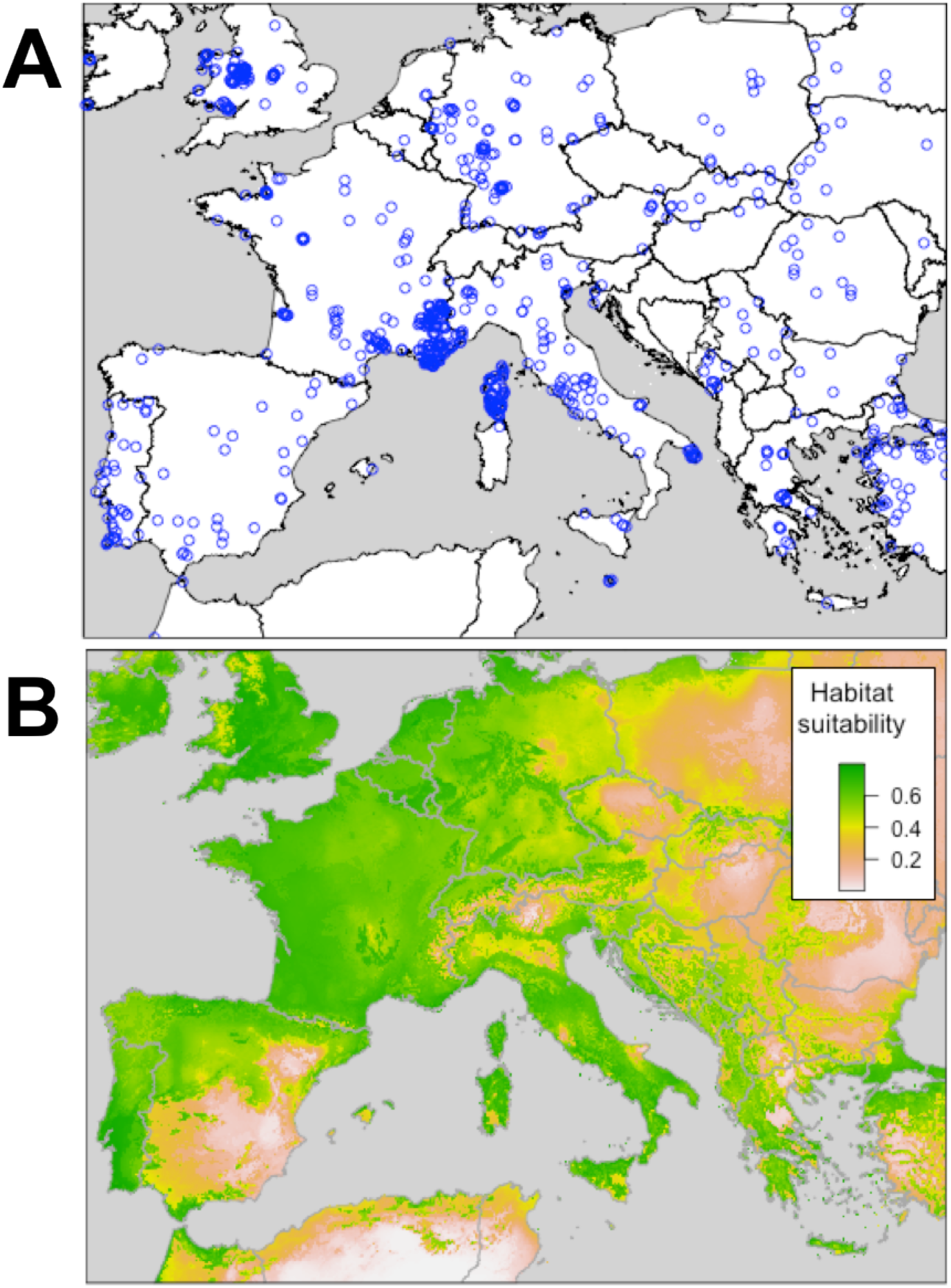
Occurrences and predicted distribution of *P. spumarius* in Europe. A. Plot of the occurrences recorded from GBIF, the literature and our own observations, B. Habitat suitability corresponds to the logistic output of the Maxent model (from red/low to green/high suitability).

## DISCUSSION

The idea of using insect as spies for the early detection of *Xf* in buffer zones and symptom-less areas is not new (Yaseen et al., 2015; D’onghia, 2017; Yaseen et al., 2017). Here, we tried to go one step further than what is done on olive groves in Southern Italy and propose to test whether insects could be used to detect, monitor or predict the distribution of *Xf*. We also propose to make a large-scale preliminary identification of the subspecies / strains of *Xf that* may be present in the ecosystem. If needed, whole genome sequencing could refine the identification.

Importantly, our study reveals some limitation of the qPCR approach sensu Harper et al. (2010 erratum 2013) to detect *Xf* in the insect vectors. The approach appears not sensitive enough to detect low bacterial load, which questions its use for the early detection of the bacterium in insects. Indeed, if we consider a result as positive only when two replicates of qPCR are positive, *Xf* will be considered as present in only two of the eleven populations and only in October. When a more sensitive approach was used no significant differences in the proportion of insects carrying *Xf* in June and in October was observed. Furthermore, the rate of undetermined results obtained with the qPCR approach is high (73.0% of the replicates with at least one undetermined result), which is unsatisfactory when it comes to the detection of plant pathogens. Although we have not formally tested a loop-mediated isothermal amplification (LAMP) approach for the detection of *Xf* in the insects (an approach increasingly used on the field), our study indicates that the results obtained with this approach should be interpreted with caution. Indeed, LAMP has been shown to be less sensitive than qPCR (Harper et al., 2010 erratum 2013; Baldi and La Porta, 2017).

This result suggests that the lower prevalence of *Xf* in *P. spumarius* observed in early season (winter-sping, 12.6%) (Yaseen et al., 2015) as compared to late season (October-December, 40%) (Elbeaino et al., 2014) in an olive grove in Italy may be artefactual. This difference may be due to the relatively poor ability of PCR and LAMP to detect *Xf* in insects with the lowest *Xf* load (more frequent in June). Consequently, we strongly advocate the use of highly sensitive methods to monitor *Xf* within insects, especially in *Xf-free* areas to avoid false negative results.

The nested PCR approach targeting *holC* optimized for the purpose of this study appeared much more sensitive than the qPCR approach and allows a first assessment of the diversity of the strains present in the environment. With this approach, all insect populations appeared to carry *Xf*, which shows that the bacterium is widely distributed in Corsica. The sampling sites of the 11 populations of *P. spumarius* tested positive for the presence of *Xf* all fall within the predicted potential distribution of the bacterium, which validates the plausibility of our nested PCR results and shows that molecular tests on insects could be used for risk assessment. It is noteworthy that while they were not visible when we performed sample collection in 2016, leaf scorch symptoms could be clearly observed in all localities tested for the presence of *Xf* when we went back to the field in October 2017. However, weather data indicated that the summer of 2017 has been the driest in 15 years and it is acknowledged that symptoms due to *Xf* are not easy to differentiate from drought symptoms (EFSA, 2015a). Thus, observed symptoms may be due to summer drought itself. However, the possibility that at least part of these symptoms are due to *Xf* cannot be ruled out as i) all populations of *P. spumarius* were tested positive for the presence of the bacterium ii) plants were found positive to *Xf* close to certain prospected sites (Fig. 1), iii) all sites were predicted as favourable for the bacterium (Fig. 5). The mechanisms underlying the interaction of water stress and infection by / sensibility to *Xf* and a possible causal relationships between these two parameters is a constant area of research (Thorne et al., 2006; Daugherty et al., 2010; Choi et al., 2013). It is difficult to assess whether or not the severe drought may have favoured the spread of *Xf* or revealed its presence. Regarding the vectors, studies conducted in the US on *Homalodisca vitripennis* Germar, 1821 have shown that the insects will take longer meal and feed more frequently on fully irrigated plants, both events that favour the acquisition and transmission of *Xf* (Krugner and Backus, 2014). This led to the conclusion that even low levels of water stress may reduce the spread of *Xf* by *H. vitripennis*. However, nothing is known about the feeding behaviour of *P. spumarius* or other European insect vectors under severe drought conditions. One can hypothesize that probing behaviour may vary with more switch from one plant to another as xylem fluid tension is reduced in all plant species. However, our sampling campaigns show that *P. spumarius* may rarely switch to woody plants. Furthermore the spittlebug is subject to aestivation. Consequently the role of other potential vectors in the spread of the disease should be investigated in the future (especially cicadas).

The wide distribution of two subspecies of *Xf* in Corsica highlighted by our molecular tests suggests that the introductions of the bacterium to Corsica may be ancient and multiple. Indeed, it appears unlikely that the bacterium spread into insect populations all over Corsica in such a short time lapse since the first detection (less than 2 years). Another argument in favour of an ancient / multiple introduction of *Xf to* Corsica is the presence of several STs, including variants either highlighted on plants (Denancé et al., 2017) and/or on insects (this study) and co-occurrence of strains / subspecies in the same matrix (plant / insect). Thus, *Polygala myrtifolia*, on which *Xf multiplex* was detected during the summer 2015, might not have been a key actor in the spread of *Xf*. This detection might just have served as a trigger for large-scale surveillance and studies that now reveal a much more complex situation than expected. Notably, the co-occurrence of subspecies / strains in the same host plant raises doubt about which entity produced symptoms and therefore on subspecies / strain occurrences used for risk assessment. Furthermore, as co-occurrence of subspecies / strains in the same host insect or plant may favour recombination, and, as a consequence enlarge host range, disease management may be further complicated (Nunney et al., 2014a; Nunney et al., 2014b; Kandel et al., 2017).

Our results indicate that the number of bacterial cells in the cibarium of *P. spumarius* may be low, even in the late season, which complicates molecular detection. Thus, our results may still underestimate the prevalence of *Xf* in insect populations. This low amount of bacteria makes possible the PCR amplification and sequencing of all the loci included in the MLST of *Xf* only on the insects in which the bacterial load is the highest. Progress should be made to circumvent this issue and capture or other approaches that are more sensitive than PCR and nested PCR may be soon implemented. *Xf multiplex* was recovered both in plants and in insects. However, we highlighted rare or widespread subspecies / variants not yet detected in plants. To the contrary, we did not detect *Xf pauca* ST53 in the targeted insect populations. These results may be explained by the followings: i) *Xf pauca* or our rare variants may be restricted to some areas where insect / plant prospection was not yet conducted, ii) some strains are harder to detect in plants (competition between PCR primers or differences in development). With survey effort, results on plants and insects should become consistent.

Obviously, we did not aim to study the entire community of vectors of *Xf* occurring in Corsica. Our goal was to gather huge populations of the same species of vector in each locality to perform molecular tests. Doing this, *P. spumarius*, which is, by the way, the only efficient vector known in Europe so far (Saponari et al., 2014; Cornara et al., 2016), was identified as the perfect candidate. While we did not aim at studying the exact phenology and host preferences of *P. spumarius*, we made a couple of biological observations that will need to be formally tested but seem nevertheless relevant to propose a few immediate solutions to decrease its potential impact on cultivated plants.

Interestingly, specimens of *P. spumarius* were easily and almost exclusively collected from *Cistus monspeliensis* with only a few specimens collected in grasses and clover. In 2016, adults started to emerge in early June, were impossible to collect in summer as mentioned by Chauvel et al. (2015) and huge adult populations reappeared in early October with the first rains. No survey was conducted in winter. We are thus unable to state whether or not adults of *P. spumarius* may survive winter in some areas. The mix of larvae and tenerals observed in many localities in early June suggest they may not survive winter but this needs to be confirmed especially in a context of global warming. Our observations contrast with observations made in Apulian olive groves (Southern Italy) by Ben Moussa et al. (2016) where *P. spumarius* begins to molt in april-may, is abundant in summer and seems more polyphagous as it moves from herbaceous plants to olive trees. It is noteworthy that *P. spumarius* is also polyphagous in Southern France as larvae were found on *ca* 120 different species of plants in the area of Montpellier (JCS, *pers. obs*.). Therefore, the situation in Corsica appears different from what has been observed elsewhere in Europe. While host preferences, exact phenology and tolerance to temperature variation of the Corsican populations of *P. spumarius* require to be precisely assessed in the future, performing a genetic analysis to evaluate their status appears also relevant. On a more general note, the contrasted observations on *P. spumarius* suggest that strategies that would need to be set up to monitor the spread of the disease would differ from areas to areas. Our field samplings and observations of huge numbers of spitlike foam on *Cistus monspeliensis* in the spring suggest that this widespread plant may play a critical role in the spread of the bacterium in Corsica. *Xf* was detected in young and older adults of *P. spumarius* which suggests that the insects could acquire the bacterium from *C. monspeliensis*, which may act as reservoir for the next season. Thus, *C. monspeliensis*, which, from our observation, seems mostlys asymptomatic to *Xf* (but was already tested positive to *Xf* on plants (Denancé et al., 2017)), could have favoured an initially invisible spread of the disease throughout Corsica. *Cistus monspeliensis* may thus have played the role of the hidden compartment suggested by one of the model selected by Soubeyrand et al. (submitted, https://www.efsa.europa.eu/sites/default/files/event/171113/171113-7.5_Soubeyrand.pdf) to best explain current observations on plants. If a key role of *P. spumarius* or *C. monspeliensis* is confirmed by further studies, disease management in Corsica may be trickier in natural ecosystems than in agro-ecosystems. Indeed, *Cistus* spp. are largely distributed in anthropized and open semi-natural habitats. They are major components of the spontaneous natural reforesting that generally follows abandonment of agriculture and grazing and are among the first colonizers after a fire. While it seems feasible to remove *Cistus* spp. from the vicinity of crops, other strategies should be implemented to control the spread of *Xf* in natural environments.

Interestingly, the species distribution model of *P. spumarius* at the European scale indicates that it may be the perfect sentinel to detect the presence of *Xf* and make a preliminary assessment of the subspecies / strains present in the environment. As a conclusion, we suggest that a study of this type in Europe will provide a better picture of the spread of the bacterium and set up a global strategy to control it. There is an urgent need to take stock of the current situation with a large-scale, blind survey and using effective / sensitive enough molecular tools. This may allow finding out why the current epidemic appears so recent, understand what could be the triggers, better design management strategies, and avoid the unnecessary economic pressure on certain geographical areas and agricultural sectors. It is even more urgent as global warming may favour the (re)-emergence of *Xf* and predictions at the European scale suggest that *Xf* may be more widespread that what is currently thought (Godefroid et al., submitted, see comments section below for preprint DOI)

## Author’s contribution

Designed the study: AC, JYR; collected samples: AC, JYR, MG, JCS, JMT; developed / wrote protocols and conducted laboratory work: AAG, SS; performed species distribution modelling: MG, JPR; discuss the results: AC, AAG, SN, JPR, SS, JYR; wrote the manuscript: AC, JYR. All authors commented on the manuscript.

## Foundings

This work was funded by grants from the SPE department of the INRA (National Agronomic Institute), the territorial collectivity of Corsica and the Horizon 2020 XF-ACTORS Project SFS-09-2016. The funders had no role in study design, data collection and analysis, decision to publish, or preparation of the manuscript.

## Acknowledgements

We thank B. Legendre from the ANSES LSV (Angers, France) for providing an inactivated bacterial suspension to test the sensitivity of our methods. We thank M.-A. Jacques (INRA, IRHS, Angers, France) for helpful discussion as well as F. Casabianca (INRA, Corte, France) and L. Hugot (CNBC Corse, France) for their collaboration in setting up a research project in Corsica. We are indebted to C. Lannou (INRA, SPE, France) for his interest and support during the course of the project.

## References

Almeida, R.P., Blua, M.J., Lopes, J.R.S., Purcell, A.H., 2005. Vector transmission of *Xylella fastidiosa:* Applying fundamental knowledge to generate disease management strategies. Annals of the Entomological Society of America 98, 775–786.

Almeida, R.P., Nunney, L., 2015. How do plant diseases caused by *Xylella fastidiosa* emerge? Plant Disease 99, 1457–1467.

ANSES, 2015. Détection de Xylella fastidiosa par PCR en temps réel sur plantes hôtes. ANSES/LSV/MA 039 version 1. Available from: https://http://www.anses.fr/fr/system/files/ANSES_MA039_Xylellafastidiosa_final.pdf

Baldi, P., La Porta, N., 2017. Xylella fastidiosa: host range and advance in molecular identification techniques. Frontiers in Plant Science 8, 944.

Ben Moussa, I.E., Mazzoni, V., Valentini, F., Yaseen, T., Lorusso, D., Speranza, S., Digiaro, M., Varvaro, L., Krügner, R., D’Onghia, A.M., 2016. Seasonal fluctuations of sap-feeding insect species infected by Xylella fastidiosa in Puglia olive groves of Southern Italy. Journal of Economic Entomology 109, 1512–1518.

Bextine, B., Tuan, S.J., Shaikh, H., Blua, M., Miller, T.A., 2004. Evaluation of methods for extracting *Xylella fastidiosa* DNA from the glassy-winged sharpshooter. Journal of Economic Entomology 97, 757–763.

Boncristiani, H., Li, J., Evans, J., Pettis, J., Chen, Y., 2011. Scientific note on PCR inhibitors in the compound eyes of honey bees, *Apis mellifera*. Apidologie 42, 457–460.

Brady, J.A., Faske, J.B., Castañeda-Gill, J.M., King, J.L., Mitchell, F.L., 2011. High-throughput DNA isolation method for detection of *Xylella fastidiosa* in plant and insect samples. Journal of Microbiological Methods 86, 310–312.

Broennimann, O., Di Cola, V., Guisan, A., 2016. ecospat: Spatial Ecology Miscellaneous Methods. version 2.1.1. https://CRAN.R-project.org/package=ecospat.

Busby, J.R., 1991. BIOCLIM: a bioclimate analysis and prediction system. Plant Protection Quarterly 6, 8–9.

Carpenter, G., Gillison, A.N., Winter, J., 1993. DOMAIN: a flexible modelling procedure for mapping potential distributions of plants and animals. Biodiversity and conservation 2, 667–680.

Chatterjee, S., Almeida, R.P.P., Lindow, S., 2008. Living in two worlds: the plant and insect lifestyles of *Xylella Fastidiosa*. Annual Review of Phytopathology 46, 243–271.

Chauvel, G., Cruaud, A., Legendre, B., Germain, J.-F., Rasplus, J.-Y., 2015. Rapport de mission d’expertise sur *Xylella fastidiosa* en Corse. Available at http://agriculture.gouv.fr/sites/minagri/files/20150908_rapport_mission_corse_xylella_31082015b.pdf.

Choi, H.-K., Iandolino, A., Silva, F.G., Cook, D.R., 2013. Water deficit modulates the response of Vitis vinifera to the Pierce’s disease pathogen *Xylella fastidiosa*. Molecular Plant-Microbe Interactions 26, 643–657.

Cornara, D., Cavalieri, V., Dongiovanni, C., Altamura, G., Palmisano, F., Bosco, D., Porcelli, F., Almeida, R.P.P., Saponari, M., 2016. Transmission of *Xylella fastidiosa* by naturally infected *Philaenus spumarius* (Hemiptera, Aphrophoridae) to different host plants. Journal of Applied Entomology 141, 80–87.

D’onghia, A.M., 2017. CIHEAM/IAMB innovative tools for early surveillance and detection of *Xylella fastidiosa*. In: D’Onghia, A.M., Brunel, S., Valentini, F. (Eds.), Xylella fastidiosa & the Olive Quick Decline Syndrome (OQDS) A serious worldwide challenge for the safeguard of olive trees - IAM Bari: CIHEAM (Centre International de Hautes Etudes Agronomiques Méditerranéennes), 2017–172 p. (Série A Mediterranean Seminars, N° 121, Options Méditerranéennes).

Daugherty, M.P., Lopes, J.R.S., Almeida, R.P.P., 2010. Strain-specific alfalfa water stress induced by *Xylella fastidiosa*. European Journal of Plant Pathology 127, 333–340.

Dellapé, G., Paradell, S., Semorile, L., Delfederico, L., 2016. Potential vectors of *Xylella fastidiosa:* a study of leafhoppers and treehoppers in citrus agroecosystems affected by Citrus Variegated Chlorosis. Entomol. Exp. Appl. 161, 92–103.

Denancé, N., Legendre, B., Briand, M., Olivier, V., de Boisseson, F., Poliakoff, F., Jacques, M.-A., 2017. Several subspecies and sequence types are associated with the emergence of Xylella fastidiosa in natural settings in France. Plant Pathology 66, 1054–1064.

EFSA, 2015a. Panel on Plant Health (PLH), European Food Safety Authority (EFSA), Scientific Opinion on the risk to plant health posed by *Xylella fastidiosa* in the EU territory, with the identification and evaluation of risk reduction, EFSA Journal 2015;13(1):3989.

EFSA, 2015b. Scientific opinion on the risks to plant health posed by Xylella fastidiosa in the EU territory, with the identification, and evaluation of risk reduction options. EFSA Journal 13, 2989.

Elbeaino, T., Yaseen, T., Valentini, F., Ben Moussa, I.E., Mazzoni, V., D’onghia, A.M., 2014. Identification of three potential insect vectors of *Xylella fastidiosa* in southern Italy. Phytopathologia Mediterranea 53, 328–332.

Elith, J., Graham, C.H., Anderson, R.P., DudÌk, M., Ferrier, S., Guisan, A., Hijmans, R.J., Huettmann, F., Leathwick, J.R., Lehmann, A., 2006. Novel methods improve prediction of species’ distributions from occurrence data. Ecography 29, 129–151.

European Plant Protection Organization, 2016. EPPO Standards PM 7 – Diagnostics PM 7/24 (2) Xylella fastidiosa. Bulletin OEPP/EPPO Bulletin 46, 463–500.

Fick, S.E., Hijmans, R.J., 2017. WorldClim 2: new 1-km spatial resolution climate surfaces for global land areas. International Journal of Climatology 37, 4302–4315.

Fielding, A.H., Bell, J.F., 1997. A review of methods for the assessment of prediction errors in conservation presence/absence models. Environmental conservation 24, 38–49.

Giampetruzzi, A., Loconsole, G., Boscia, D., Calzolari, A., Chiumenti, M., Martelli, G.P., Saldarelli, P., Almeida, R.P., Saponari, M., 2015. Draft genome sequence of CO33, a coffee-infecting isolate of *Xylella fastidiosa*. Genome Announcements 3, e01472–01415.

Gu, X., 1995. Maximum likelihood estimation of the heterogeneity of substitution rate among nucleotide sites. Molecular Biology and Evolution 12, 546–557.

Harper, S.J., Ward, L.I., Clover, G.R.G., 2010 erratum 2013. Development of LAMP and real-time PCR methods for the rapid detection of Xylella fastidiosa for quarantine and field applications. Phytopathology 12, 1282–1288.

Hijmans, R.J., E., C.S., Parra, J.L., Jones, P.G., Jarvis, A., 2005. Very high resolution interpolated climate surfaces for global land areas. International Journal of Climatology 25, 1965–1978.

Hijmans, R.J., Phillips, S., Leathwick, J., Elith, J., 2016. dismo: Species Distribution Modeling. R package version 1.1-4. https://CRAN.R_project.org/package=dismo.

Hirzel, A.H., Le Lay, G., Helfer, V., Randin, C., Guisan, A., 2006. Evaluating the ability of habitat suitability models to predict species presences. Ecological Modelling 199, 142–152.

Huson, D.H., Bryant, D., 2006. Application of phylogenetic networks in evolutionary studies. Molecular Biology and Evolution 23, 254–267.

Jacques, M.-A., Denancé, N., Legendre, B., Morel, E., Briand, M., Mississipi, S., Durand, K., Olivier, V., Portier, P., Poliakoff, F., Crouzillat, D., 2016. New coffee-infecting Xylella fastidiosa variants derived via homologous recombination. Applied and Environmental Microbiology 82, 1556–1568.

Janse, J., Obradovic, A., 2010. *Xylella fastidiosa:* its biology, diagnosis, control and risks. Journal of Plant Pathology, S35–S48.

Jiménez-Valverde, A., Peterson, A.T., Soberón, J., Overton, J.M., Aragón, P., Lobo, J.M., 2011. Use of niche models in invasive species risk assessments. Biological Invasions 13, 2785–2797.

Kandel, P.P., Almeida, R.P., Cobine, P.A., De La Fuente, L., 2017. Natural competence rates are variable among *Xylella fastidiosa* strains and homologous recombination occurs in vitro between subspecies *fastidiosa* and *multiplex*. Molecular Plant-Microbe Interactions 30, 589–600.

Krell, R.K., Boyd, E.A., Nay, J.E., Park, Y.L., Perring, T.M., 2007. Mechanical and insect transmission of *Xylella fastidiosa* to *Vitis vinifera*. American Journal of Enology and Viticulture 58, 211–216.

Krugner, R., Backus, E.A., 2014. Plant water stress effects on stylet probing behaviors of *Homalodisca vitripennis* (Hemiptera: Cicadellidae) associated with acquisition and inoculation of the bacterium *Xylella fastidiosa*. Journal of Economic Entomology 107, 66–74.

Liu, C., Berry, P.M., Dawson, T.P., Pearson, R.G., 2005a. Selecting thresholds of occurrence in the prediction of species distributions. Ecography 28, 385–393.

Liu, C., Berry, P.M., Dawson, T.P., Pearson, R.G., 2005b. Selecting thresholds of occurrence in the prediction of species distributions. 28, 385–393.

Loconsole, G., Saponari, M., Boscia, D., D’Attoma, G., Morelli, M., Martelli, G.P., Almeida, R.P.P., 2016. Intercepted isolates of Xylella fastidiosa in Europe reveal novel genetic diversity. European Journal of Plant Pathology 146, 85.

Merow, C., Smith, M.J., Silander, J.A., 2013. A practical guide to MaxEnt for modeling species’distributions: what it does, and why inputs and settings matter. Ecography 36, 1058–1069.

Muscarella, R., Galante, P.J., Soley-Guardia, M., Boria, R.A., Kass, J.M., Uriarte, M., Anderson, R.P., 2014. ENMeval: An R package for conducting spatially independent evaluations and estimating optimal model complexity forMaxent ecological niche models. Methods in Ecology and Evolution 5, 1198–1205.

Nunney, L., Elfekih, S., Stouthamer, R., 2012a. The importance of multilocus sequence typing: Cautionary tales from the bacterium *Xylella fastidiosa*. Phytopathology 102, 456–460.

Nunney, L., Hopkins, D.L., Morano, L.D., Russell, S.E., Stouthamer, R., 2014a. Intersubspecific recombination in *Xylella fastidiosa* strains native to the United States: Infection of novel hosts associated with an unsuccessful invasion. Applied and Environmental Microbiology 80, 1159–1169.

Nunney, L., Schuenzel, E.L., Scally, M., Bromley, R.E., Stouthamer, R., 2014b. Large-scale intersubspecific recombination in the plant-pathogenic bacterium *Xylella fastidiosa* is associated with the host shift to mulberry. Applied and Environmental Microbiology 80, 3025–3033.

Nunney, L., Vickerman, D.B., Bromley, R.E., Russell, S.A., Hartman, J.R., Morano, L.D., Stouthamer, R., 2013. Recent evolutionary radiation and host plant specialization in the *Xylella fastidiosa* subspecies native to the United States. Applied and Environmental Microbiology 79, 2189–2200.

Nunney, L., Yuan, X., Bromley, R.E., Stouthamer, R., 2012b. Detecting genetic introgression: high levels of intersubspecific recombination found in *Xylella fastidiosa* in Brazil. Applied and Environmental Microbiology 78, 4702–4714.

Olmo, D., Nieto, A., Adrover, F., Urbano, A., Beidas, O., Juan, A., Marco-Noales, E., López, M.M., Navarro, I., Monterde, A., Montes-Borrego, M., Navas-Cortés, J.A., Landa, B.B., 2017. First detection of Xylella fastidiosa infecting cherry (Prunus avium) and Polygala myrtifolia plants, in Mallorca Island, Spain. Plant Disease 101, 1820–1820.

Paião, F., Meneguim, A., Casagrande, E., Leite, R., 2002. Envolvimento de cigarras (Homoptera, Cicadidae) na transmissão de *Xylella fastidiosa* em cafeeiro. Fitopatol. Bras. 27, 67.

Peterson, A.T., Nakazawa, Y., 2008. Environmental data sets matter in ecological niche modelling: an example with *Solenopsis invicta* and *Solenopsis richteri*. Global Ecology and Biogeography 17, 135–144.

Phillips, S.J., Anderson, R.P., Schapire, R.E., 2006. Maximum entropy modeling of species geographic distributions. Ecological Modelling 190, 231–259.

Phillips, S.J., Dudik, M., 2008. Modeling of species distributions with Maxent: new extensions and a comprehensive evaluation. Ecography 31, 161–175.

Purcell, A.H., 2013. Paradigms: examples from the bacterium *Xylella fastidiosa*. Annual Review of Phytopathology 51, 339–356.

Purcell, A.H., Finlay, A., 1979. Evidence for non-circulative transmission of Pierce’s disease bacterium by sharpshooter leafhoppers. Phytopathology 69, 393–395.

Radosavljevic, A., Anderson, R.P., 2014. Making better Maxent models of species distributions: complexity, overfitting and evaluation. Journal of Biogeography 41, 629–643.

Rambaut, A., 2006. FigTree. Available online at http://tree.bio.ed.ac.uk/software/figtree/.

Redak, R.A., Purcell, A.H., Lopes, J.R., Blua, M.J., Mizel, R.F.r., Andersen, P.C., 2004a. The biology of Xylem fluid-feeding insect vectors of *Xylella fastidiosa* and their relation to disease epidemiology. Annual Review of Entomology 49, 243–270.

Redak, R.A., Purcell, A.H., Lopes, J.R.S., Blua, M.J., Mizell, R.F., Andersen, P.C., 2004b. The biology of xylem fluid-feeding insect vectors of *Xylella fastidiosa* and their relation to disease epidemiology. Annu. Rev. Entomol. 49, 243–270.

Retchless, A.C., Labroussaa, F., Shapiro, L., Stenger, D.C., Lindow, S.E., Almeida, R.P., 2014. Genomic insights into *Xylella fastidiosa* interactions with plant and insect hosts. In: Heidelberg, S.B. (Ed.), Genomics of Plant-Associated Bacteria pp. 177–202.

Saponari, M., Boscia, D., Nigro, F., Martelli, G.P., 2013a. Identification of DNA sequences related to *Xylella fastidiosa* in oleander, almond and olive trees exhibiting leaf scorch symptoms in Apulia (Southern Italy). Journal of Plant Pathology 95, 668.

Saponari, M., Loconsole, G., Cornara, D., Yokomi, R.K., De Stradis, A., Boscia, D., Bosco, D., Martelli, G.P., Krugner, R., Porticelli, F., 2014. Infectivity and transmission of Xylella fastidiosa by Philaenus spumarius (Hemiptera: Aphrophoridae) in Apulia, Italy. Journal of Economic Entomology 107, 1316–1319.

Saponari, M., Loconsole, G., Liao, H.H., Jiang, B., Savino, V., Yokomi, R.K., 2013b. Validation of high-throughput real time polymerase chain reaction assays for simultaneous detection of invasive citrus pathogens. J. Virol. Methods 193, 478–486.

Schrader, C., Schielke, A., Ellerbroek, L., Johne, R., 2012. PCR inhibitors - occurrence, properties and removal. Journal of Applied Microbiology 113, 1014–1026.

Severin, H.H.P., 1949. Transmission of the virus of Pierce’s disease by leafhoppers. Hilgardia 19, 190–202.

Severin, H.H.P., 1950. Spittle-insect vectors of Pierce’s disease virus. II. Life history and virus transmission. Hilgardia 19, 357–382.

Shamim, G., Ranjan, S.K., Pandey, D.M., Ramani, R., 2014. Biochemistry and biosynthesis of insect pigments. European Journal of Entomology 111, 149–164.

Shcheglovitova, M., Anderson, R.P., 2013. Estimating optimal complexity for ecological niche models: a jackknife approach for species with small sample sizes. Ecological Modelling 269, 9–17.

Stamatakis, A., 2006. Phylogenetic models of rate heterogeneity: A High Performance Computing Perspective. International Parallel and Distributed Processing Symposium (IPDPS 2006), Rhodes Island, Greece, 8 pp.

Stamatakis, A., 2014. RAxML version 8: a tool for phylogenetic analysis and post-analysis of large phylogenies. Bioinformatics 30, 1312–1313.

Thorne, E.T., Stevenson, J.F., Rost, T.L., Labavitch, J.M., Matthews, M.A., 2006. Pierce’s disease symptoms: comparison with symptoms of water deficit and the impact of water deficits. American Journal of Enology and Viticulture 57, 1–11.

Yang, Z., 1994. Maximum likelihood phylogenetic estimation from DNA sequences with variable rates over sites: approximate methods. Journal of Molecular Evolution 39, 306–314.

Yaseen, T., Drago, S., Valentini, F., Elbeaino, T., Stampone, G., Digiaro, M., D’Onghia, A.M., 2015. On site detection of *Xylella fastidiosa* in host plants and in “spy insects” using the real-time loop-mediated isothermal amplification method. Phytopathologia Mediterranea 54, 488–496.

Yaseen, T., Valentini, F., Santoro, F., Lorusso, D., D’Onghia, A.M., 2017. The “Spy Insect” approach for monitoring *Xylella fastidiosa* in absence of symptomatic plants In: D’Onghia, A.M., Brunel, S., Valentini, F. (Eds.), Xylella fastidiosa & the Olive Quick Decline Syndrome (OQDS) A serious worldwide challenge for the safeguard of olive trees - IAM Bari: CIHEAM (Centre International de Hautes Etudes Agronomiques Méditerranéennes), 2017 – 172 p. (Série A Mediterranean Seminars, N° 121, Options Méditerranéennes).

Yuan, X., Morano, L., Bromley, R., Spring-Pearson, S., Stouthamer, R., Nunney, L., 2010. Multilocus sequence typing of *Xylella fastidiosa* causing Pierce’s disease and oleander leaf scorch in the United States. Phytopathology 100, 601–611.

